# *In silico* prioritization of transporter-drug relationships from drug sensitivity screens

**DOI:** 10.1101/381335

**Authors:** A. César-Razquin, E. Girardi, M. Yang, M. Brehme, J. Sáez-Rodríguez, G. Superti-Furga

**Affiliations:** CeMM Research Center for Molecular Medicine of the Austrian Academy of Sciences, Vienna, Austria; Heidelberg University, Heidelberg, Germany; Joint Research Centre for Computational Biomedicine (JRC-COMBINE), RWTH Aachen University, Faculty of Medicine, Aachen, Germany; European Molecular Biology Laboratory - European Bioinformatics Institute, Wellcome Genome Campus, Cambridge, UK; CBmed - Center for Biomarker Research in Medicine GmbH, Graz, Austria; Center for Physiology and Pharmacology, Medical University of Vienna, Vienna, Austria

**Keywords:** Solute carriers (SLCs), ABC transporters (ATP binding cassette), drug sensitivity and resistance, drug transport, regularized linear regression

## Abstract

The interplay between drugs and cell metabolism is a key factor in determining both compound potency and toxicity. In particular, how and to what extent transmembrane transporters affect drug uptake and disposition is currently only partially understood. Most transporter proteins belong to two protein families: the ATP-Binding Cassette (ABC) transporter family, whose members are often involved in xenobiotic efflux and drug resistance, and the large and heterogeneous family of Solute carriers (SLCs). We recently argued that SLCs are collectively a rather neglected gene group, with most of its members still poorly characterized, and thus likely to include many yet-to-be-discovered associations with drugs. We searched publicly available resources and literature to define the currently known set of drugs transported by ABCs or SLCs, which involved ~500 drugs and more than 100 transporters. In order to extend this set, we then mined the largest publicly available pharmacogenomics dataset, which involves approximately 1000 molecularly annotated cancer cell lines and their response to 265 chemical compounds, and used regularized linear regression models (Elastic Net, LASSO) to predict drug responses based on SLC and ABC data (expression levels, SNVs, CNVs). The most predictive models included both known and previously unidentified associations between drugs and transporters. To our knowledge, this represents the first application of regularized linear regression to this set of genes, providing an extensive prioritization of potentially pharmacologically interesting interactions.

## Introduction

The role of cellular metabolism in determining the potency and distribution of drugs is increasingly recognized (Zhao et al., 2013). Along with the enzymes involved in actual xenobiotic transformation, such as members of the cytochrome and transferases families, a critical role is played by transmembrane transporters, which directly affect both the uptake and the excretion of drugs and their metabolites (Zhou et al., 2017). Among transmembrane transporters, two large families have been described: the family of ABC (ATP-binding cassette) transporters and the family of Solute carriers (SLCs) (Hediger et al., 2013). ABC transporters are pumps powered by the hydrolysis of ATP and show a remarkable broad range of substrates, including lipids, secondary metabolites and xenobiotics. Members of this family, such as the ABCB/MDR and ABCC/MRP proteins, have been associated with resistance to a large number of structurally diverse compounds in cancer cells (Fletcher et al., 2010). Solute carriers (SLCs) are secondary transporters involved in uptake or efflux of metabolites and other chemical matter (Cesar-Razquin et al., 2015). At more than 400 members and counting, SLCs represent the second largest family of membrane proteins and comprise uniporters, symporters and antiporters, further grouped into subfamilies based on sequence similarity (Hoglund et al., 2011). Among the reported SLC substrates are all major building blocks of the cell, such as nucleic acids, sugars, lipids and aminoacids as well as vitamins, metals and other ions (Hediger et al., 2013). Despite the critical processes mediated by these proteins, a large portion of SLCs is still poorly characterized and, in several cases, lacks any associations with a substrate (Cesar-Razquin et al., 2015). Importantly, several members of the SLCO (also known as Organic Anion Transporter Proteins or OATPs) and SLC22 families (including the group of organic cation transporters or OCTs, organic zwitterion/cation transporters or OCTNs and organic anion transporters or OATs) have been found to play prominent roles in the uptake and excretion of drugs, especially in the liver and kidneys (Hagenbuch and Stieger, 2013). Several other cases of Solute carriers mediating the uptake of drugs have been reported, such as in the case of methotrexate and related anti-folate drugs with the folate transporter SLC19A1 (Zhao et al., 2011) or the anti-cancer drug YM155/sepantronium bromide and the orphan transporter SLC35F2 (Winter et al., 2014). Indeed, whether carrier-mediated uptake is the rule or rather the exception is still a matter of discussion (Dobson and Kell, 2008; Sugano et al., 2010). Due to the understudied nature of transporters and SLCs in particular, we can nonetheless expect that several other associations between drugs and transporters, involving direct transport or indirect effects, remain to be discovered and could provide novel insights into the pharmacokinetics of drugs and drug-like compounds.

Analysis of basal gene expression and genomic features in combination with drug sensitivity data allows the identification of molecular markers that render cells both sensitive and resistant to specific drugs. Such a pharmacogenomic analysis represents a powerful method to prioritize *in silico* gene-compound associations. Different statistical and machine learning (ML) strategies have been used in the past to confirm known as well as to identify novel drug-gene associations, although generally in a genome-wide context (Iorio et al., 2016). For our study, we mined the “Genomics of Drug Sensitivity in Cancer” (GDSC) dataset (Iorio et al., 2016) which contains drug sensitivity data to a set of 265 compounds over ~1000 molecularly annotated cancer cell lines, in order to explore drug relationships exclusively involving transporters (SLCs and ABCs). To such end, we used regularized linear regression (Elastic Net, LASSO) to generate predictive models from which to extract cooperative sensitivity and resistance drug-transporter relationships, in what represents, to our knowledge, the first work applying this type of analysis to this group of genes.

## Materials and Methods

### Data

SLC and ABC genes were considered as in (Cesar-Razquin et al., 2015). Known drug transport cases involving SLC and ABC proteins were obtained from four main repositories as of September 2017: DrugBank (Law et al., 2014), The IUPHAR/BPS Guide to PHARMACOLOGY (Alexander et al., 2015), KEGG: Kyoto Encyclopedia of Genes and Genomes (Kanehisa and Goto, 2000), and UCSF-FDA TransPortal (Morrissey et al., 2012). These data were complemented with various other cases found in the literature (Sprowl and Sparreboom, 2014; Winter et al., 2014; Nigam, 2015; Radic-Sarikas et al., 2017). Source files were parsed using custom python scripts, and all entries were manually curated, merged together and redundancies eliminated. The final compound list was searched against PubChem (Kim et al., 2016) in order to systematize names. *f*A list of FDA-approved drugs was obtained from the organization’s website. Network visualization was done using Cytoscape (Shannon et al., 2003).

All data corresponding to the Genomics of Drug Sensitivity in Cancer (GDSC) dataset (drug sensitivity, expression, copy number variations, single nucleotide variants, compounds, cell lines) were obtained from the original website of the project http://www.cancerrxgene.org/downloads as of September 2016. Drug sensitivity and transcriptomics data were used as provided. Genomics data were transformed into a binary matrix of genomic alterations vs cell lines, where three different modifications for every gene were considered using the original source files: amplifications (ampSLCx), deletions (delSLCx) and variants (varSLCx). An amplification was annotated if there were more than two copies of at least one of the alleles for the gene of interest, and a deletion if at least one of the alleles was missing. Single nucleotide variants were filtered in order to exclude synonymous SNVs as well as nonsynonymous SNVs predicted not to be deleterious by either SIFT (Ng and Henikoff, 2001), Polyphen2 (Adzhubei et al., 2010) or FATHMM (Shihab et al., 2013).

### LASSO regression

LASSO regression analysis was performed using the ‘glmnet’ R package (Friedman et al., 2010). Expression values for all genes in the dataset (17419 genes in total) were used as input features. For each compound, the analysis was iterated 50 times over 10-fold cross validation. At each cross validation, features were ranked based on their frequency of appearance (number of times a feature has non zero coefficient for 100 default lambda possibilities). We then averaged the ranking across the 500 runs (50 iterations x 10 CV) in order to obtain a final list of genes associated to each compound. In this context, the most predictive gene for a certain drug does not necessarily have an average rank of one, even though its final rank is first.

### Elastic Net regression

Elastic Net regression analysis was performed using the ‘glmnet’ R package (Friedman et al., 2010). Genomic data (copy number variations and single nucleotide variants) and transcriptional profiles of SLC and ABC genes across the cell line panel were used as input variables, either alone or in combination. Drug AUC values were used as response. Elastic Net parameters were fixed as follows: i) alpha, the mixing parameter that defines the penalty, was set to 0.5 in order to apply an intermediate penalty between Ridge and LASSO, and ii) lambda, the tuning parameter that controls the overall strength of the penalty, was determined individually for every model (drug) by optimizing the mean squared cross-validated error.

For each compound, 500 Elastic Net models were generated by a 100x 5-fold cross-validation procedure. In order to assess model performance, the Concordance Index (Harrell et al., 1996; Papillon-Cavanagh et al., 2013) between the predicted and observed AUC values was calculated for each run, and then averaged across all models. This index estimates the fraction of cell line pairs for which the model correctly predicts which of the two is the most and least sensitive; hence CI values of 0.5 and 1 would indicate random and perfect predictors, respectively. Feature weights were calculated by normalizing the fitted model coefficients to the absolute maximum at every cross-validation run. The final list of features associated with each compound was built by computing the frequency of appearance of each feature in all the 500 models as well as its average weight. Features with positive weights are associated with a resistance phenotype to the compound, and negative weights are indicative of sensitivity.

## Results

### SLC and ABCs as drug transporters

We collected data from public repositories as well as relevant publications to define the current knowledge on transport of chemical compounds by members of the SLC and ABC protein classes. A total of 493 compounds linked to 107 transporters were retrieved, which altogether formed a single large network with a few other smaller components (**Fig.1, Table S1**).

**Figure 1.**
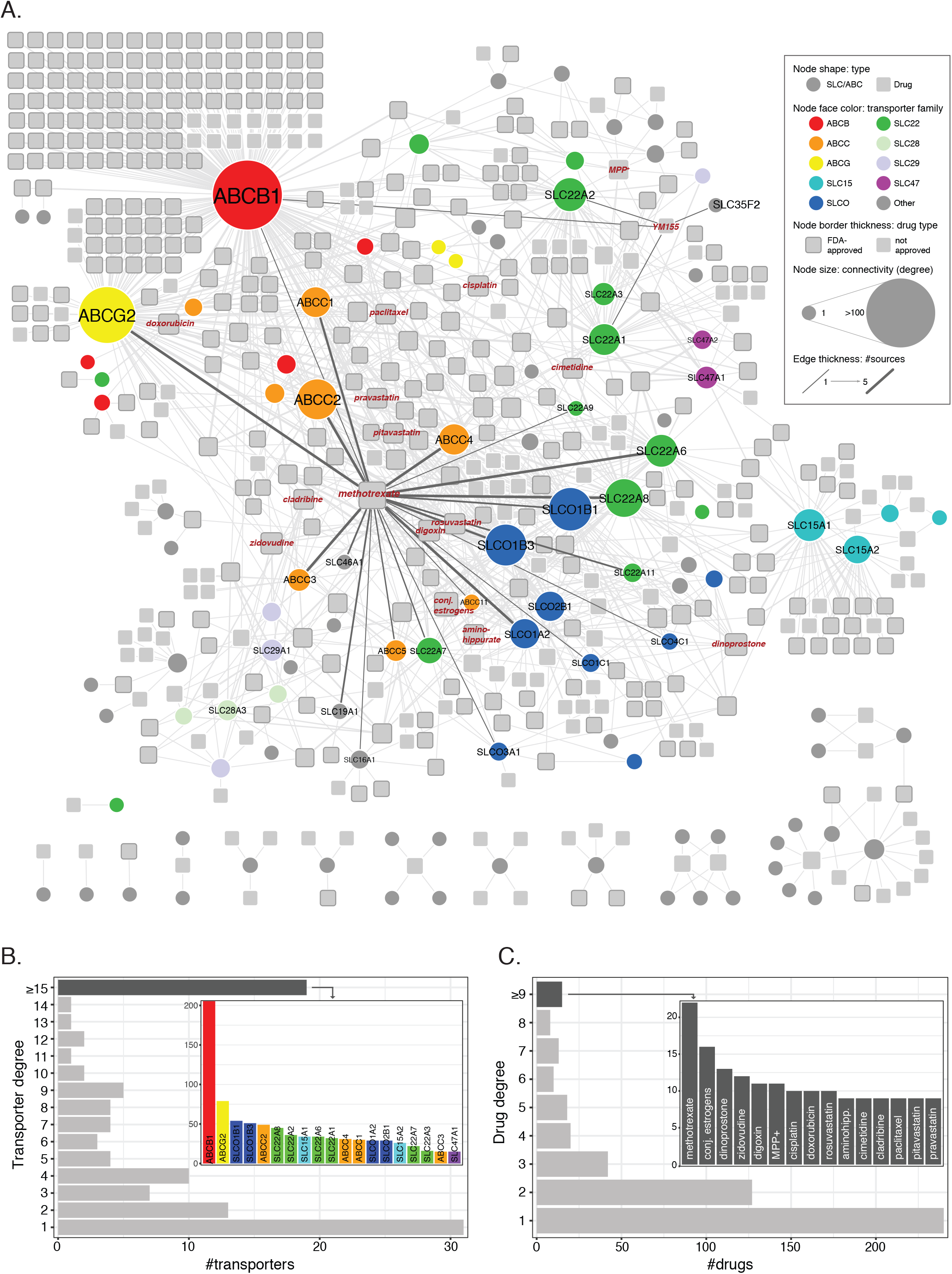
**A)** Network visualization of known SLC/ABC-mediated drug transport cases. Circular nodes represent SLC and ABC transporters, and squares represent chemical compounds. Drugs approved by the FDA (Food and Drug Administration) are displayed with thicker gray borders. Edges connect transporters to compounds and their thickness indicates the number of sources supporting each connection (see Methods). Size indicates node degree (number of edges incident to the node). Relevant transporter families are color coded. **B)** Transporter degree distribution. The inlet barchart displays the transporters connected to at least 15 compounds. Bar colors correspond to transporter families in A. **C)** Same as B for drugs.

Within the largest network and in agreement with previous reports (Nigam, 2015), three families are significantly enriched (hypergeometric test, FDR ≤ 0.05): the SLCO/SLC21 family of organic anion transporters (9/12 members) (Hagenbuch and Stieger, 2013), the SLC22 family of organic anion, cation and zwitterion transporters (13/23) (Koepsell, 2013; Nigam, 2018), and the ABCC family of multidrug resistance transporters (8/13) (Vasiliou et al., 2009). Not surprisingly, ABCB1 (P-glycoprotein; MDR1), the very well-studied efflux pump known for its broad substrate specificity and mediation of resistance to a large amount of drugs (Aller et al., 2009), is the most connected transporter in the network, linked to more than 200 compounds. In particular, 106 compounds are connected exclusively with ABCB1 and 25 other are exclusively shared with ABCG2 (BCRP), another well-known transporter and the one with the second highest degree in the network (Robey et al., 2007) (**Fig.1B**). Other top-connected SLCs include members of the above mentioned SLCO and SLC22 families, which also show several common substrates (e.g. SLCO1B1 and SLCO1B3 share 36 compounds, and SLC22A8 and SLC22A6 share 20), as well as members of the SLC15 family (SLC15A1 and SLC15A2, which share 22 compounds), involved in the transport of beta-lactam antibiotics and peptide-mimetics (Smith et al., 2013). In contrast to these cases, other transporters appear related to one or only a few compounds. One such case is SLC35F2, whose only reported substrate to date is the anti-cancer drug YM155 (sepantronium bromide) (Winter et al., 2014). Finally, while most chemical compounds appear linked to one or two transporters, a few others show higher connectivities (**Fig.1C**). A well-known example, methotrexate is transported by more than 20 different SLC and ABCs, including some belonging to families not commonly involved in drug transport, such as the folate carriers SLC19A1 and SLC46A1.

### Transporter expression landscape in cancer cell lines

The GDSC dataset contains expression data for 371 SLCs and 46 ABCs across a panel of ~1000 cell lines of different tissue origin. Each of these cell lines effectively express between 167 and 255 transporters, with a median value of 195 (**Fig.2A**). Although all together they cover almost the whole transporter repertoire (414/417), the distribution is clearly bimodal, with a common set of ~130 transporters expressed in at least 900 cell lines and a more specific set of ~140 expressed in less than 100 (**Fig.2B**). Among the most commonly expressed transporters, we find several members of the SLC25 (mitochondrial carriers) and SLC35 (nucleoside-sugars transporters) sub-families, the two largest among SLCs, as well as several members of the SLC39 family of zinc transporters. On the other end, many members of the SLC22 family, a large and well known group of proteins involved in the transport of drugs, as well as the SLC6 family, a well-studied family of neurotransmitter transporters, show a more specific expression pattern. As for ABCs, it is worth highlighting that subfamilies A and C present half of their members in the set of transporters of specific expression, while subfamily B has members in both sets.

**Figure 2.**
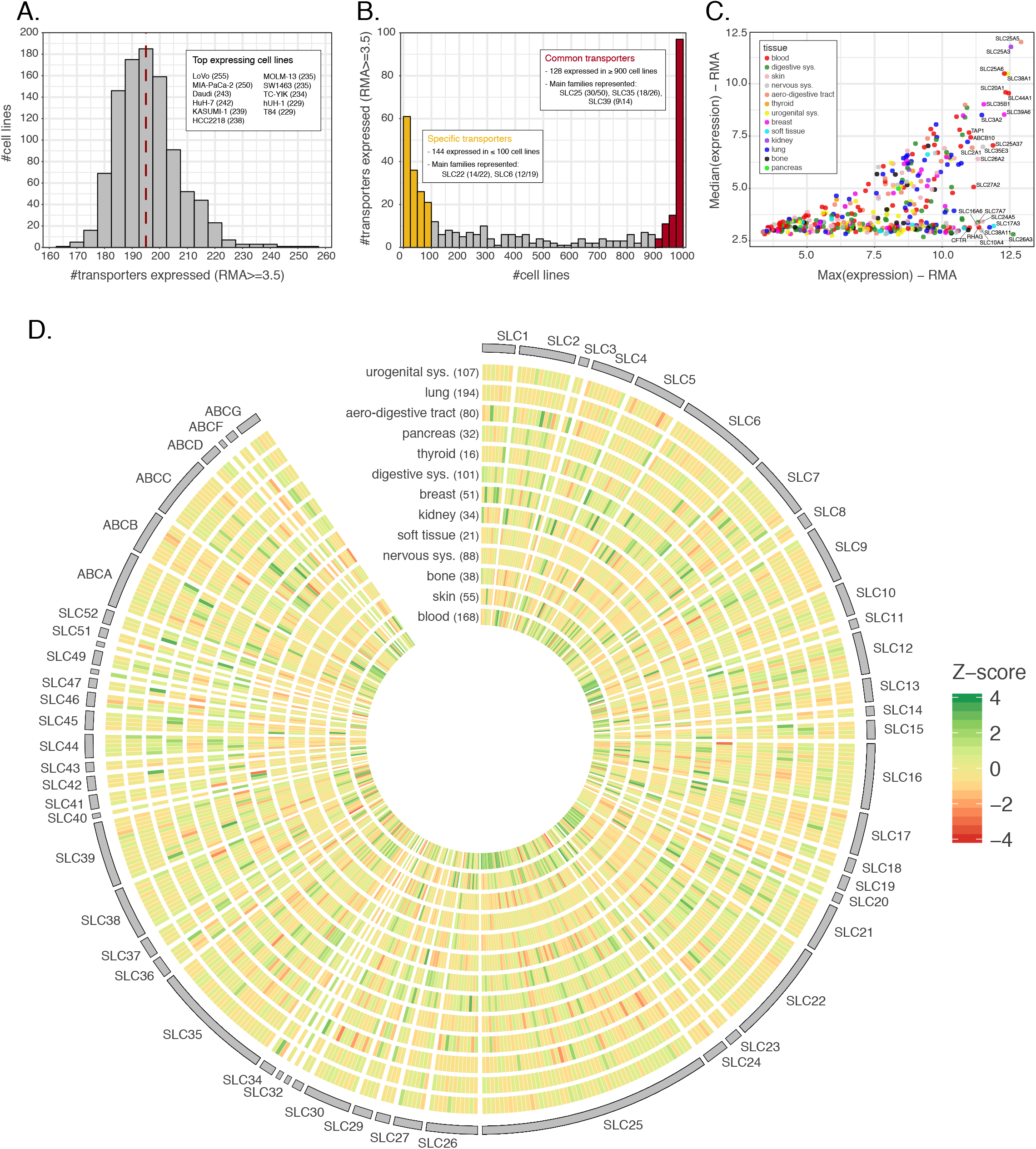
**A)** Number of transporters (SLCs and ABCs) expressed across cell lines in GDSC dataset. A cut-off of 3.5 in RMA units is set to consider a gene as expressed (~73% genes expressed). The red line indicates the median number of transporters expressed per cell line. The inlet lists the 11 cell lines expressing the highest number of transporters, indicated between parentheses. **B)** Number of cell lines expressing each of the transporters. The color bars and inlets indicate sets of transporters showing more common or specific expression across the panel. **C)** Median expression vs maximum expression for each transporter across the cell line panel. Color indicates the tissue of origin of the cell line presenting the maximum expression for the transporter. **D)** Transporter Z-scores of the average expression values within each tissue. Tissue names with number of cell lines between parenthesis are indicated on the x-axis. Transporters are ordered alphabetically by family and name.

When looking at actual expression values across the panel, some of the commonly expressed transporters coincide with those of highest expression (**Fig.2C**). The most extreme cases are SLC25A5, SLC25A3, SLC25A6 and SLC38A1, which present very similar maximum and median values across the cell line panel. On the contrary, other transporters such as SLC26A3, SLC17A3, or SLC38A11 present a much wider range of expression, being amongst the highest expressed in some cell lines but completely absent from others.

Finally, substantial differences become apparent when considering transporter expression patterns according to the tissue of origin of the GDSC cancer cell lines (**Fig.2D**), Cell lines belonging to the hematopoietic (blood) lineage, which includes leukemias, lymphomas and myelomas, present the largest proportion of transporters with highest average expression values (28%), as indicated by Z-score, followed by cancer cell lines belonging to skin, kidney and the digestive system. This indicates a broad spectrum of transporters being present in cell lines of these tissue origins. Interestingly, kidney cell lines also present the largest number of transporters with low expression values, pointing to a very wide range of expression and high specificity in those cells.

### LASSO regression shows importance of SLC genes across whole genome

We investigated the importance of SLC and ABC transporters for drug response by applying regularized linear regression on the GDSC dataset. To this end, we first built LASSO models of sensitivity to each compound based on genome-wide gene expression levels (17419 genes in total) (Tibshirani, 1996), and then looked for cases where a transporter ranked as the top (first) predictor (see Methods). The choice of the LASSO method is motivated by its ability to shrink a large number of coefficients to zero, ideal for models that make use of thousands of predictors. Moreover, being a linear regression method, it can account for both positive and negative interactions (i.e. resistance and sensitivity, for example by export and import in the case of a transporter), thus increasing the interpretability of the results. The decision to focus exclusively on the top predictor is supported by a literature search. Indeed, the average number of PubMed publications containing both the drug and the gene name was over 40 in the case of top predictors, falling down to below 10 for the ones ranked second (**Fig.S1**).

Consistent with their well-characterized role as drug-transporters, the multi-drug resistance pump ABCB1, as well as ABCG2, were the main predictors of resistance to a large number of drugs (**Table 1A**). More interestingly, several compounds had an SLC as their best predictor (**Table 1B**). Among them, and in concordance with previous expression-sensitivity data (Rees et al., 2016), we find the sensitive association of sepantronium bromide (YM155) and SLC35F2, its main known importer (Winter et al., 2014). Another sensitive association involving SLC35F2 links this transporter to NSC-207895, a MDMX inhibitor (Wang et al., 2011). DMOG (dimethyloxalylglycin), a synthetic analogue of a-ketoglutarate that inhibits HIF prolyl hydroxylase (Zhdanov et al., 2015), showed association to two SLCs: monocarboxylate transporter SLC16A7 (MCT2) was the main predictor for sensitivity to this compound, while creatine transporter SLC6A8 (CT1) was associated with resistance. However, due to the high IC50 values of DMOG (in the millimolar range), this association is unlikely to be clinically relevant. Finally, cystine-glutamate transporter SLC7A11 (Blomen et al., 2015) is associated to resistance to the ROS-inducing drugs Shikonin, (5Z)-7-Oxozeaenol and Piperlongumine. This is in agreement with previous studies that highlighted a positive correlation of the expression of this transporter and resistance to several drugs via import of the cystine necessary for glutathione balance maintenance (Huang et al., 2005).

**Table 1A.** LASSO ABC-drug top associations.

| LASSO Top Hits, all<br>17419 genes used | Top sensitive associations<br>(average rank) | Top resistant associations<br>(average rank) |
| --- | --- | --- |
| **ABCB1** |  | YM155 (1)<br>Paclitaxel (1.1)<br>BI-2536 (6.0)<br>A-443654 (32)<br>Vinorelbine (1)<br>Thapsigargin (20)<br>AT-7519 (1.8)<br>WZ3105 (1)<br>PHA-793887 (2.2)<br>GSK690693 (15)<br>Vinblastine (1.1)<br>Docetaxel (1.2)<br>ZM447439 (77)<br>ZG-10 (1.3)<br>QL-VIII-58 (1)<br>QL-XII-61 (9.7) |
| **ABCG2** |  | CUDC-101 (12)<br>THZ-2-102-1 (1.8) |
| ABCA10 | STF-62247 (20)<br>FR-180204 (22) |  |

**Table 1B.** LASSO SLC-drug top associations.

| LASSO Top Hits, all<br>17419 genes used | Top sensitive associations<br>(average rank) | Top resistant associations<br>(average rank) |
| --- | --- | --- |
| SLC16A7 | DMOG (1) |  |
| SLC6A8 |  | DMOG (40) |
| SLC30A2 |  | CP724714 (28) |
| SLC35F2 | YM155 (2.24) |  |
| SLC35F2 | NSC-207895 (9.5) |  |
| SLC7A11 |  | Shikonin (2) |
| SLC7A11 |  | (5Z)-7-Oxozeaenol (12) |
| SLC7A11 |  | piperlongumine (12) |

### Elastic Net regression identifies transporter-drug relationships

In order to further explore SLC and ABC involvement in drug response, we decided to build new predictive models based on transporter molecular data only. By removing the effect of other genes in the models, we can prioritize compounds that show a stronger dependency on transporters, as well as to analyze potential cooperative interactions among them. Given the smaller amount of predictors in this case, we used Elastic Net regression, a generalization of the LASSO that overcomes some of its limitations and that has already been applied in similar contexts (Zou, 2005; Barretina et al., 2012; Iorio et al., 2016). Assessment of model performance was done by cross-validation using the Concordance Index (CI) (see Methods).

We considered different predictors to build the models: genomics (Copy Number Variations and Single Nucleotide Variants), transcriptomics (gene expression) and a combination of both. Among these, gene expression resulted to be most predictive, in agreement with previous reports (Aydin et al., 2014)(**Fig.3A**). 139 (53%) of the 265 drugs included in the dataset had predictive models with a CI higher than 0.60, and 36 (14%) higher than 0.65 (**Fig.3B**). For those drugs, we then ranked genes based on their frequency of appearance in the cross-validated models (indicative of the robustness of the association) and their average weight (indicative of the strength of the association as well as its direction). In this context increased levels of transporter expression could therefore be associated with either sensitivity or resistance to the drug, for example through its uptake or efflux, respectively (**Fig.3C**). Among the top ranked transporter-drug associations, we identified several known cases of drug transport. For instance, the strongest sensitivity association with sepantronium bromide (YM155) corresponded again to SLC35F2. Similarly, the strongest resistance association for this drug was ABCB1, which includes YM155 among its many substrates (Lamers et al., 2012; Voges et al., 2016; Radic-Sarikas et al., 2017). Another example was methotrexate, for which the folate transporter SLC19A1, known to mediate its import (Zhao et al., 2011), ranked second for sensitivity association (**Table S3**).

**Figure 3.**
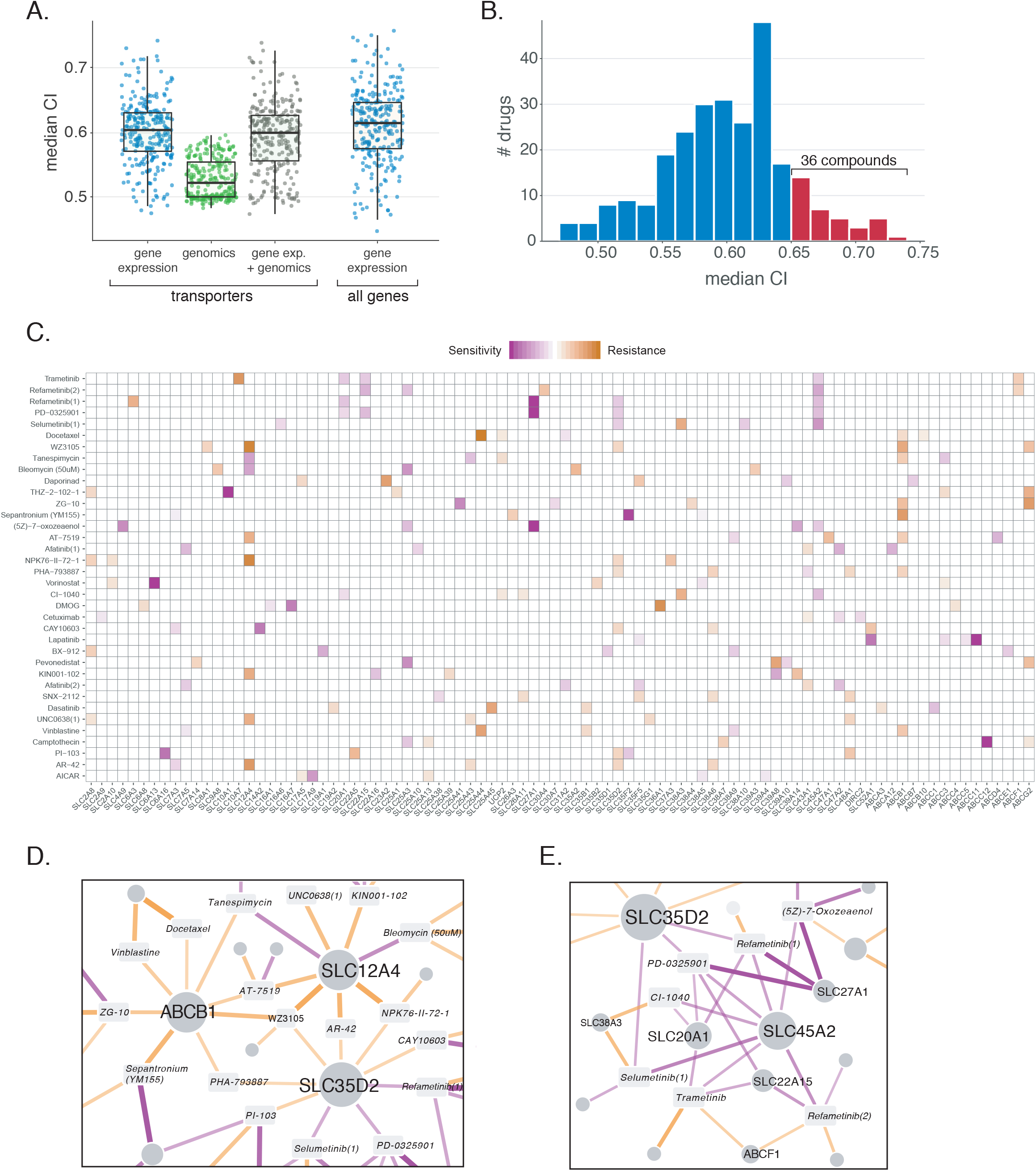
**A)** Comparison of Elastic Net regression performance (Concordance Index) using different input data: gene expression, genomics (CNVs and SNVs) and a combination of both. **B)** CI value distribution using gene expression as input. Red bars indicate drugs with a median CI higher than 0.65, which were selected for subsequent analysis. **C)** Elastic Net results for drugs with the highest CI values. The top 5 associations are shown for each compound. Purple indicates associations linked to sensitivity (higher expression value confers sensitivity to the compound), and orange indicates resistance. **E)** Network representation of three transporters appearing as “hubs” (e.g. connected to several different compounds) in the results, including the well-known multidrug resistance protein ABCB1. **D)** Same as E for MEK inhibitors, which show a similar association pattern.

Two major patterns are apparent in the set of top-ranking associations: genes showing similar profiles of resistance or sensitivity across several different and unrelated compounds as well as groups of genes showing a similar profile in relation to a functionally related class of drugs (**Fig. 3C**).

A prototypical case of the first pattern is ABCB1, which is associated with resistance phenotypes to several compounds (**Fig.3D**). Together with the aforementioned YM155, resistance relationships were predicted for known substrates vinblastine and docetaxel (Fletcher et al., 2010), 17-AAG/Tanespimycin (Huang et al., 2007) and AT-7519 (Cihalova et al., 2015) as well as other not previously associated compounds such as ZG-10 (a JNK1 inhibitor), the CDK2/5/7 inhibitor PHA-793887 and the broad kinase inhibitor WZ3105. Similar to ABCB1, other transporters showed multiple resistance and sensitivity associations to different compounds, particularly kinases and chromatin-related enzymes. Two of these “hubs” were SLC12A4/KCC1, a potassium-chloride cotransporter involved in cell volume homeostasis (Arroyo et al., 2013), and SLC35D2, an activated sugar transporter localized in the Golgi (Song, 2013).

As an example of the second class of associations, some of the best models were achieved for compounds belonging to the MEK inhibitor drug class (Trametinib, Selumetinib, Refametinib, CI-1040, PD-0325901, (5Z)-7-oxozeaenol), which showed very similar patterns, with sensitivity associated to SLC45A2, SLC27A1, SLC20A1, and SLC22A15 (**Fig.3E**). SLC45A2 has been related to melanin synthesis and it is highly expressed in melanomas (Park et al., 2017), a cancer type sensitive to MEK inhibitors. Interestingly, SLC20A1/PiT1, a sodium-dependent phosphate transporter (Olah et al., 1994), was previously shown to regulate the ERK1/2 pathway independently of phosphate transport in skeletal cells (Bon et al., 2018). SLC27A1, a long-chain fatty acid transporter, and SLC22A15, an orphan member of the well-known family of cationic transporters involved in the transport of different compounds, were not previously associated with this drug class.

Finally, we also observed a strong sensitivity relationship between expression levels of the amino acid transporter SLC7A5/LAT1 and the Her2 and EGFR kinase inhibitors Afatinib, Gefitinib and Bosutinib (**Fig. 2C**), consistent with previously published data (Timpe et al., 2015).

## Discussion

Transporters of the ABC and SLC superfamilies are increasingly recognized as key players in determining the distribution and metabolism of drugs and other xenobiotic compounds as they possess distinct and extremely variable expression patterns across cell lines and tissues (O’Hagan et al., 2018). Moreover, they have been implicated in the development of resistance to several chemotherapeutic drugs (Fletcher et al., 2010). A survey of currently known drug transport relationships revealed that only a fifth of the more than 500 SLCs and ABCs have been described to be involved in the transport of drugs. These transporters appear to be very unevenly distributed, with some genes and families considerably more represented and better connected than others (**Fig.1**). This is the case for several members of the ABCB, ABCC, SLCO and SL22 sub-families. Similarly, while compounds such as methotrexate are linked to more than 20 transporters, most drugs are connected to only one.

To further expand this network, we took advantage of the expression and drug sensitivity data available within the GDSC project. We started by characterizing the expression patterns of SLCs and ABCs in the GDSC panel of ~1000 cancer cell lines, covering thirteen different tissues of origin (**Fig.2**). Roughly 80% of SLCs and 90% of ABCs were included in the datasets and we observed a bimodal distribution of their expression, with similarly-sized sets of transporters either present in most cell lines or specific to a few. A large variability in the level of expression was also observed within the superfamilies, consistent with what recently reported by another recent study (O’Hagan et al., 2018).

We then implemented a linear regression-based approach to identify the set of transporters associated with sensitivity to each compound across all cell lines. Previous reports undertook a similar approach to identify associations of the ABC (Szakacs et al., 2004) and SLCO/SLC22 (Okabe et al., 2008) families with drug sensitivity within a limited set of about 60 cell lines. We now extended these results to a much more comprehensive set of cell lines while implementing regularized linear regression approaches (Elastic Net and LASSO regression). We identify a large set of drug-transporter associations roughly split between sensitivity and resistance relationships (**Tables 1A and 1B, Fig.3**). Known associations involving, for example, ABCB1 expression levels with increasing resistance to several unrelated compounds as well as known interactions such as the associations between antifolates and SLC19A1 or YM155 and SLC35F2 were clearly identified. Interestingly, we also observed cases were, similarly to ABCB1, a single gene was associated with several compounds, possibly as a result of an alteration of the general metabolic state of the cell. We also observed the opposite scenario, with several genes associated with a functionally related class of compounds as in the case of the MEK inhibitors and the genes SLC45A2, SLC27A1, SLC20A1, and SLC22A15. To our knowledge, no transporter has so far been identified for any member of this class of compounds, and while the association with the skin-specific SLC45A2 transporter is likely the result of the high sensitivity of melanoma cell lines to these drugs, other associations are more difficult to interpret.

We propose the gene list reported here as a means of prioritizing transporters that could explain the transport and pharmacodynamics of the associated compounds. While in many cases these associations could be due to indirect effects, such as a change in the metabolic state of the cells that renders them more sensitive or resistant to a compound, some might correspond to actual import or export processes. Further validation, for example modulating the expression levels of the transporters or by transport assays, will be necessary in order to confirm and distinguish such different scenarios. Finally, the power of the analysis could also be increased by larger datasets, for instance including additional compounds, as well as by orthogonal or more accurate measurements. Availability of such pharmacogenomics datasets will be of critical importance for the further identification and characterization of transporter-drug associations. In conclusion, we provide here an overview of the known ABC- and SLC-based drug transport relationships and expand this with an *in* silico-derived ranking of transporter-drug associations, identifying several novel and potential interesting interactions that could affect the pharmacodynamics and pharmacokinetics of a large set of clinically relevant compounds.

## Acknowledgments

CeMM and the Superti-Furga laboratory are supported by the Austrian Academy of Sciences (G.S.-F. and A.C.R.). We acknowledge receipt of third-party funds from the Austrian Science Fund (FWF I2192-B22 ERASE, A.C.R and FWF P29250-B30 VITRA, E.G), and from the JRC for Computational Biomedicine which was partially funded by Bayer AG.

## Author contributions

ACR, MY performed the data analysis. EG, MB, JSR and GSF provided scientific insight and project supervision. ACR, EG, MY, JSR and GSF wrote the manuscript.

---

**Supplemental Figure 1.**
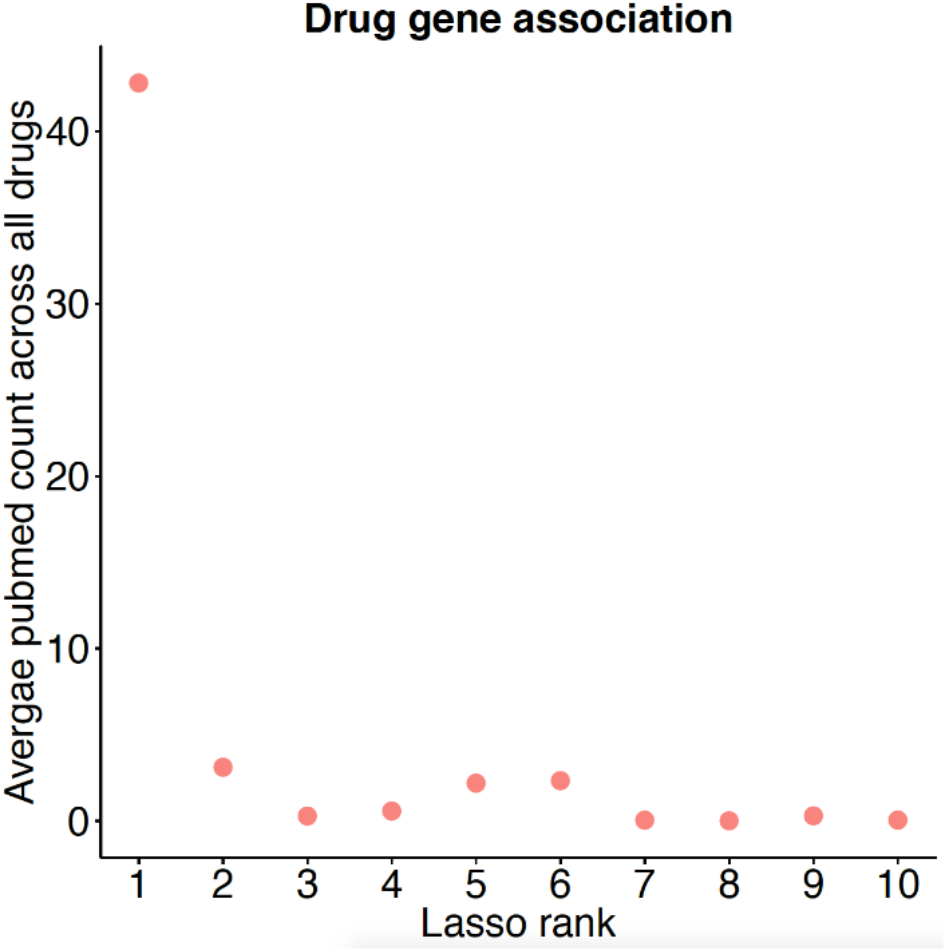
PubMed search of drug gene associations.

## References

Adzhubei, I.A., Schmidt, S., Peshkin, L., Ramensky, V.E., Gerasimova, A., Bork, P., et al. (2010). A method and server for predicting damaging missense mutations. Nat Methods 7(4), 248–249. doi: 10.1038/nmeth0410-248.

Alexander, S.P., Kelly, E., Marrion, N., Peters, J.A., Benson, H.E., Faccenda, E., et al. (2015). The Concise Guide to PHARMACOLOGY 2015/16: Transporters. Br J Pharmacol 172(24), 6110–6202. doi: 10.1111/bph.13355.

Aller, S.G., Yu, J., Ward, A., Weng, Y., Chittaboina, S., Zhuo, R., et al. (2009). Structure of P-glycoprotein reveals a molecular basis for poly-specific drug binding. Science 323(5922), 1718–1722. doi: 10.1126/science.1168750.

Arroyo, J.P., Kahle, K.T., and Gamba, G. (2013). The SLC12 family of electroneutral cation-coupled chloride cotransporters. Mol Aspects Med 34(2-3), 288–298. doi: 10.1016/j.mam.2012.05.002.

Aydin, I., Weber, S., Snijder, B., Samperio Ventayol, P., Kuhbacher, A., Becker, M., et al. (2014). Large scale RNAi reveals the requirement of nuclear envelope breakdown for nuclear import of human papillomaviruses. PLoS Pathog 10(5), e1004162. doi: 10.1371/journal.ppat.1004162.

Barretina, J., Caponigro, G., Stransky, N., Venkatesan, K., Margolin, A.A., Kim, S., et al. (2012). The Cancer Cell Line Encyclopedia enables predictive modelling of anticancer drug sensitivity. Nature 483(7391), 603–607. doi: 10.1038/nature11003.

Blomen, V.A., Majek, P., Jae, L.T., Bigenzahn, J.W., Nieuwenhuis, J., Staring, J., et al. (2015). Gene essentiality and synthetic lethality in haploid human cells. Science 350(6264), 1092–1096. doi: 10.1126/science.aac7557.

Bon, N., Couasnay, G., Bourgine, A., Sourice, S., Beck-Cormier, S., Guicheux, J., et al. (2018). Phosphate (Pi)-regulated heterodimerization of the high-affinity sodium-dependent Pi transporters PiT1/Slc20a1 and PiT2/Slc20a2 underlies extracellular Pi sensing independently of Pi uptake. J Biol Chem 293(6), 2102–2114. doi: 10.1074/jbc.M117.807339.

Cesar-Razquin, A., Snijder, B., Frappier-Brinton, T., Isserlin, R., Gyimesi, G., Bai, X., et al. (2015). A Call for Systematic Research on Solute Carriers. Cell 162(3), 478–487. doi: 10.1016/j.cell.2015.07.022.

Cihalova, D., Staud, F., and Ceckova, M. (2015). Interactions of cyclin-dependent kinase inhibitors AT-7519, flavopiridol and SNS-032 with ABCB1, ABCG2 and ABCC1 transporters and their potential to overcome multidrug resistance in vitro. Cancer Chemother Pharmacol 76(1), 105–116. doi: 10.1007/s00280-015-2772-1.

Dobson, P.D., and Kell, D.B. (2008). Carrier-mediated cellular uptake of pharmaceutical drugs: an exception or the rule? Nat Rev Drug Discov 7(3), 205–220. doi: 10.1038/nrd2438.

Fletcher, J.I., Haber, M., Henderson, M.J., and Norris, M.D. (2010). ABC transporters in cancer: more than just drug efflux pumps. Nat Rev Cancer 10(2), 147–156. doi: 10.1038/nrc2789.

Friedman, J., Hastie, T., and Tibshirani, R. (2010). Regularization Paths for Generalized Linear Models via Coordinate Descent. J Stat Softw 33(1), 1–22.

Hagenbuch, B., and Stieger, B. (2013). The SLCO (former SLC21) superfamily of transporters. Mol Aspects Med 34(2-3), 396–412. doi: 10.1016/j.mam.2012.10.009.

Harrell, F.E., Jr., Lee, K.L., and Mark, D.B. (1996). Multivariable prognostic models: issues in developing models, evaluating assumptions and adequacy, and measuring and reducing errors. Stat Med 15(4), 361–387. doi: 10.1002/(SICI)1097-0258(19960229)15:4<361::AID-SIM168>3.0.CO;2-4.

Hediger, M.A., Clemencon, B., Burrier, R.E., and Bruford, E.A. (2013). The ABCs of membrane transporters in health and disease (SLC series): introduction. Mol Aspects Med 34(2-3), 95–107. doi: 10.1016/j.mam.2012.12.009.

Hoglund, P.J., Nordstrom, K.J., Schioth, H.B., and Fredriksson, R. (2011). The solute carrier families have a remarkably long evolutionary history with the majority of the human families present before divergence of Bilaterian species. Mol Biol Evol 28(4), 1531–1541. doi: 10.1093/molbev/msq350.

Huang, Y., Blower, P.E., Liu, R., Dai, Z., Pham, A.N., Moon, H., et al. (2007). Chemogenomic analysis identifies geldanamycins as substrates and inhibitors of ABCB1. Pharm Res 24(9), 1702–1712. doi: 10.1007/s11095-007-9300-x.

Huang, Y., Dai, Z., Barbacioru, C., and Sadee, W. (2005). Cystine-glutamate transporter SLC7A11 in cancer chemosensitivity and chemoresistance. Cancer Res 65(16), 7446–7454. doi: 10.1158/0008-5472.CAN-04-4267.

Iorio, F., Knijnenburg, T.A., Vis, D.J., Bignell, G.R., Menden, M.P., Schubert, M., et al. (2016). A Landscape of Pharmacogenomic Interactions in Cancer. Cell 166(3), 740–754. doi: 10.1016/j.cell.2016.06.017.

Kanehisa, M., and Goto, S. (2000). KEGG: kyoto encyclopedia of genes and genomes. Nucleic Acids Res 28(1), 27–30.

Kim, S., Thiessen, P.A., Bolton, E.E., Chen, J., Fu, G., Gindulyte, A., et al. (2016). PubChem Substance and Compound databases. Nucleic Acids Res 44(D1), D1202–1213. doi: 10.1093/nar/gkv951.

Koepsell, H. (2013). The SLC22 family with transporters of organic cations, anions and zwitterions. Mol Aspects Med 34(2-3), 413–435. doi: 10.1016/j.mam.2012.10.010.

Lamers, F., Schild, L., Koster, J., Versteeg, R., Caron, H.N., and Molenaar, J.J. (2012). Targeted BIRC5 silencing using YM155 causes cell death in neuroblastoma cells with low ABCB1 expression. Eur J Cancer 48(5), 763–771. doi: 10.1016/j.ejca.2011.10.012.

Law, V., Knox, C., Djoumbou, Y., Jewison, T., Guo, A.C., Liu, Y., et al. (2014). DrugBank 4.0: shedding new light on drug metabolism. Nucleic Acids Res 42(Database issue), D1091–1097. doi: 10.1093/nar/gkt1068.

Morrissey, K.M., Wen, C.C., Johns, S.J., Zhang, L., Huang, S.M., and Giacomini, K.M. (2012). The UCSF-FDA TransPortal: a public drug transporter database. Clin Pharmacol Ther 92(5), 545–546. doi: 10.1038/clpt.2012.44.

Ng, P.C., and Henikoff, S. (2001). Predicting deleterious amino acid substitutions. Genome Res 11(5), 863–874. doi: 10.1101/gr.176601.

Nigam, S.K. (2015). What do drug transporters really do? Nat Rev Drug Discov 14(1), 29–44. doi: 10.1038/nrd4461.

Nigam, S.K. (2018). The SLC22 Transporter Family: A Paradigm for the Impact of Drug Transporters on Metabolic Pathways, Signaling, and Disease. Annu Rev Pharmacol Toxicol 58, 663–687. doi: 10.1146/annurev-pharmtox-010617-052713.

O’Hagan, S., Wright Muelas, M., Day, P.J., Lundberg, E., and Kell, D.B. (2018). GeneGini: Assessment via the Gini Coefficient of Reference “Housekeeping” Genes and Diverse Human Transporter Expression Profiles. Cell Syst 6(2), 230–244 e231. doi: 10.1016/j.cels.2018.01.003.

Okabe, M., Szakacs, G., Reimers, M.A., Suzuki, T., Hall, M.D., Abe, T., et al. (2008). Profiling SLCO and SLC22 genes in the NCI-60 cancer cell lines to identify drug uptake transporters. Mol Cancer Ther 7(9), 3081–3091. doi: 10.1158/1535-7163.MCT-08-0539.

Olah, Z., Lehel, C., Anderson, W.B., Eiden, M.V., and Wilson, C.A. (1994). The cellular receptor for gibbon ape leukemia virus is a novel high affinity sodium-dependent phosphate transporter. J Biol Chem 269(41), 25426–25431.

Papillon-Cavanagh, S., De Jay, N., Hachem, N., Olsen, C., Bontempi, G., Aerts, H.J., et al. (2013). Comparison and validation of genomic predictors for anticancer drug sensitivity. J Am Med Inform Assoc 20(4), 597–602. doi: 10.1136/amiajnl-2012-001442.

Park, J., Talukder, A.H., Lim, S.A., Kim, K., Pan, K., Melendez, B., et al. (2017). SLC45A2: A Melanoma Antigen with High Tumor Selectivity and Reduced Potential for Autoimmune Toxicity. Cancer Immunol Res 5(8), 618–629. doi: 10.1158/2326-6066.CIR-17-0051.

Radic-Sarikas, B., Halasz, M., Huber, K.V.M., Winter, G.E., Tsafou, K.P., Papamarkou, T., et al. (2017). Lapatinib potentiates cytotoxicity of YM155 in neuroblastoma via inhibition of the ABCB1 efflux transporter. Sci Rep 7(1), 3091. doi: 10.1038/s41598-017-03129-6.

Rees, M.G., Seashore-Ludlow, B., Cheah, J.H., Adams, D.J., Price, E.V., Gill, S., et al. (2016). Correlating chemical sensitivity and basal gene expression reveals mechanism of action. Nat Chem Biol 12(2), 109–116. doi: 10.1038/nchembio.1986.

Robey, R.W., Polgar, O., Deeken, J., To, K.W., and Bates, S.E. (2007). ABCG2: determining its relevance in clinical drug resistance. Cancer Metastasis Rev 26(1), 39–57. doi: 10.1007/s10555-007-9042-6.

Shannon, P., Markiel, A., Ozier, O., Baliga, N.S., Wang, J.T., Ramage, D., et al. (2003). Cytoscape: a software environment for integrated models of biomolecular interaction networks. Genome Res 13(11), 2498–2504. doi: 10.1101/gr.1239303.

Shihab, H.A., Gough, J., Cooper, D.N., Stenson, P.D., Barker, G.L., Edwards, K.J., et al. (2013). Predicting the functional, molecular, and phenotypic consequences of amino acid substitutions using hidden Markov models. Hum Mutat 34(1), 57–65. doi: 10.1002/humu.22225.

Smith, D.E., Clemencon, B., and Hediger, M.A. (2013). Proton-coupled oligopeptide transporter family SLC15: physiological, pharmacological and pathological implications. Mol Aspects Med 34(2-3), 323–336. doi: 10.1016/j.mam.2012.11.003.

Song, Z. (2013). Roles of the nucleotide sugar transporters (SLC35 family) in health and disease. Mol Aspects Med 34(2-3), 590–600. doi: 10.1016/j.mam.2012.12.004.

Sprowl, J.A., and Sparreboom, A. (2014). Uptake carriers and oncology drug safety. Drug Metab Dispos 42(4), 611–622. doi: 10.1124/dmd.113.055806.

Sugano, K., Kansy, M., Artursson, P., Avdeef, A., Bendels, S., Di, L., et al. (2010). Coexistence of passive and carrier-mediated processes in drug transport. Nat Rev Drug Discov 9(8), 597–614. doi: 10.1038/nrd3187.

Szakacs, G., Annereau, J.P., Lababidi, S., Shankavaram, U., Arciello, A., Bussey, K.J., et al. (2004). Predicting drug sensitivity and resistance: profiling ABC transporter genes in cancer cells. Cancer Cell 6(2), 129–137. doi: 10.1016/j.ccr.2004.06.026.

Tibshirani, R. (1996). Regression Shrinkage and Selection via the Lasso. Journal of the Royal Statistical Society (Series B) 58, 267–288.

Timpe, L.C., Li, D., Yen, T.Y., Wong, J., Yen, R., Macher, B.A., et al. (2015). Mining the Breast Cancer Proteome for Predictors of Drug Sensitivity. J Proteomics Bioinform 8(9), 204–211. doi: 10.4172/jpb.1000370.

Vasiliou, V., Vasiliou, K., and Nebert, D.W. (2009). Human ATP-binding cassette (ABC) transporter family. Hum Genomics 3(3), 281–290.

Voges, Y., Michaelis, M., Rothweiler, F., Schaller, T., Schneider, C., Politt, K., et al. (2016). Effects of YM155 on survivin levels and viability in neuroblastoma cells with acquired drug resistance. Cell Death Dis 7(10), e2410. doi: 10.1038/cddis.2016.257.

Wang, H., Ma, X., Ren, S., Buolamwini, J.K., and Yan, C. (2011). A small-molecule inhibitor of MDMX activates p53 and induces apoptosis. Mol Cancer Ther 10(1), 69–79. doi: 10.1158/1535-7163.MCT-10-0581.

Winter, G.E., Radic, B., Mayor-Ruiz, C., Blomen, V.A., Trefzer, C., Kandasamy, R.K., et al. (2014). The solute carrier SLC35F2 enables YM155-mediated DNA damage toxicity. Nat Chem Biol 10(9), 768–773. doi: 10.1038/nchembio.1590.

Zhao, R., Diop-Bove, N., Visentin, M., and Goldman, I.D. (2011). Mechanisms of membrane transport of folates into cells and across epithelia. Annu Rev Nutr 31, 177–201. doi: 10.1146/annurev-nutr-072610-145133.

Zhao, Y., Butler, E.B., and Tan, M. (2013). Targeting cellular metabolism to improve cancer therapeutics. Cell Death Dis 4, e532. doi: 10.1038/cddis.2013.60.

Zhdanov, A.V., Okkelman, I.A., Collins, F.W., Melgar, S., and Papkovsky, D.B. (2015). A novel effect of DMOG on cell metabolism: direct inhibition of mitochondrial function precedes HIF target gene expression. Biochim Biophys Acta 1847(10), 1254–1266. doi: 10.1016/j.bbabio.2015.06.016.

Zhou, F., Zhu, L., Wang, K., and Murray, M. (2017). Recent advance in the pharmacogenomics of human Solute Carrier Transporters (SLCs) in drug disposition. Adv Drug Deliv Rev 116, 21–36. doi: 10.1016/j.addr.2016.06.004.

Zou, H.H., T. (2005). Regularization and variable selection via the elastic net. Journal of the Royal Statistical Society (Series B) 67(2), 301–320.

